# Nitrate-Reducing Commensals Reshape Oral Biofilm Ecology and Reveal Hcp as a Critical Determinant of *Porphyromonas gingivalis* Persistence

**DOI:** 10.64898/2026.08.06.743060

**Authors:** B. Ross Belvin, Janina P. Lewis

## Abstract

Dietary nitrate (NO₃⁻) supplementation is emerging as a promising strategy for suppressing oral pathobionts through microbial generation of reactive nitrogen species (RNS), including nitrite (NO₂⁻) and nitric oxide (NO). However, the mechanisms that enable periodontal pathogens to survive nitrate-derived nitrosative stress within polymicrobial communities remain poorly understood. Previously, we identified the hybrid cluster protein (Hcp) as a major nitrosative stress defense factor in *Porphyromonas gingivalis* demonstrating ∼170-fold induction of *hcp* expression following nitrite exposure and as a requirement for survival at physiologically relevant nitrite concentrations. Here we investigated the role of Hcp in promoting *P. gingivalis* persistence within nitrate-reducing biofilms. Using human *ex vivo* plaque biofilms, we found that Hcp is essential for *P. gingivalis* survival under both basal and nitrate-supplemented conditions. In a defined nine-species biofilm model, nitrate reduction suppressed wild-type *P. gingivalis*, whereas deletion of *hcp* (*Δhcp*) resulted in complete population clearance. Metatranscriptomics revealed that nitrate-induced *hcp* expression was not restricted to *P. gingivalis* but was part of a coordinated nitrosative stress response shared among oral anaerobes, including *Prevotella intermedia*, *Fusobacterium nucleatum*, and *Veillonella atypica*. Moreover, nitrate reduction disrupted a previously synergistic interaction between *Veillonella spp.* and *P. gingivalis*, converting a supportive relationship into an inhibitory microenvironment that constrained pathogen survival. Collectively, these findings identify Hcp-mediated nitrosative stress resistance as a major determinant of fitness within nitrate-reducing biofilms and reveal RNS as key ecological force shaping interactions between commensal nitrate reducers and periodontal pathogens. These results provide a mechanistic framework linking dietary nitrate metabolism to oral microbiome homeostasis.

## Introduction

Periodontitis is a chronic inflammatory disease initiated by microbial biofilms that accumulate on tooth surfaces and trigger a persistent host immune response^1^. Because these complex biofilm communities cannot be efficiently eliminated, inflammation becomes sustained, resulting in destruction of gingival tissues, alveolar bone loss, and ultimately tooth loss^2^. Beyond the oral cavity, periodontitis is strongly associated with systemic conditions including rheumatoid arthritis, diabetes, and cardiovascular diseases^3–5^.

A defining feature of periodontitis is the transition from a symbiotic to a dysbiotic oral microbiome. This shift is characterized by the loss of beneficial commensals and an expansion of pro-inflammatory anaerobic pathobionts, including the keystone pathogen *Porphyromonas gingivalis*^6–8^. Through its arsenal of virulence factors, *P. gingivalis* disrupts host mechanisms and promotes the growth of other disease-associated species, driving chronic inflammation and tissue destruction^6,9,10^.

Recent studies highlight the importance of oral inorganic nitrate (NO_3_) in maintaining oral microbiome homeostasis^11^. Salivary glands concentrate plasma nitrate in the oral cavity resulting in the highest concentrations found in the human body, and is then reduced to nitrite (NO_2_) by oral commensals such as *Veillonella, Rothia,* and *Neisseria* ^12,13,14^. This NO_3_ →NO_2_ → NO enterosalivary circuit elevates oral and systemic nitrite and nitric oxide (NO) promoting cardiovascular health^15–17^. This simultaneously shapes the oral microbiome by selecting for nitrate reducers and taxa that can survive the elevated levels of nitrosative stress^18^. Consequently, inorganic nitrate supplementation has emerged as a promising prebiotic strategy to suppress dysbiotic pathogens while promoting symbiotic communities^19^. Clinical trials demonstrate nitrate rich diets promote systemic health and a health associated oral microbiome^20–24^. *Ex vivo* biofilm studies using subgingival plaque or salivary microbes demonstrate an increase in health associated microbes and repression of disease associated microbes^19,25,26^.

As an obligate anaerobe with extensive iron dependent metabolism, *P. gingivalis* is particularly susceptible to nitrosative stress because NO and related reactive nitrogen species readily target iron-sulfur proteins^27^. *P. gingivalis* utilizes the heme dependent transcription regulator HcpR to sense nanomolar concentrations of NO and regulate the expression of *hcp*^28–30^. Hcp is a hybrid cluster protein that acts as a putative high affinity NO reductase^31,32^. Hcp is ubiquitously found among facultative and obligate Gram negative anaerobes where it functions as a primary RNS detoxifier^32^. In our previous studies, *hcp* was identified as one of the most highly induced genes during nitrosative stress, exhibiting approximately 170 fold upregulation following exposure to nitrite^29^. Importantly, we demonstrated that Hcp is essential for nitrosative stress resistance: wild type *P. gingivalis* survives in the presence of 2mM NO_2_ but an *hcp* mutant fails to grow at concentrations above 0.2mM nitrite^33^. These findings established the HcpR-Hcp pathway as a critical survival mechanism in *P. gingivalis*.

These observations have important therapeutic implications. Nitrate supplementation has emerged as a promising prebiotic strategy, capable of promoting health associated microbial communities while suppressing periodontal pathogens. However, the molecular mechanisms connecting nitrate metabolism, nitrosative stress, and pathogen suppression remain poorly understood. In this study, we build on our studies from defined nitrite exposure experiments to biologically relevant polymicrobial systems in which nitrite is generated by oral commensals through dietary nitrate metabolism. Specifically, nitrate-reducing bacteria such as *Veillonella*, *Rothia*, and *Neisseria* convert dietary nitrate to nitrite, creating a nitrosative environment that is predicted to be hostile to *P. gingivalis*. Using both *ex vivo* plaque biofilms and a defined nine-species oral biofilm model, we demonstrate that Hcp is required for *P. gingivalis* survival under these conditions. Metatranscriptomic analyses further reveal that nitrate induces a coordinated community-wide response characterized by widespread *hcp* upregulation. Moreover, nitrate disrupts the cooperative interaction between *Veillonella spp*. and *P. gingivalis,* transforming a previously supportive relationship into an inhibitory microenvironment. Together, these findings identify Hcp-mediated nitrosative stress resistance as a critical determinant of pathogen survival and provide a mechanistic explanation for how dietary nitrate promotes a health-associated oral microbiome.

## Materials and Methods

Detailed experimental procedures are provided in Supplementary Methods. A schematic overview of the experimental *ex vivo* biofilm design is shown in Fig. 5A.

### Bacterial strains and growth conditions

*Porphyromonas gingivalis* W83, *Prevotella intermedia* OMA14, *Fusobacterium nucleatum* (ATCC 25586), *Capnocytophaga sputigina* (ATCC 33612), *Veillonella atypica* (KON), *Streptococcus oralis* (ATCC 9811), *Streptococcus gordonii* (ATCC 35105), *Neisseria perflava* (ATCC 14799), and *Corynebacterium matruchotii* (ATCC 33612) were grown using standard anaerobic or aerobic conditions as appropriate for each species. Biofilm experiments were performed in a protein-rich artificial saliva medium designed to mimic the oral environment. Detailed growth conditions and medium compositions are provided in Supplementary Methods.

### Construction of *P. gingivalis* mutant and complemented strains

The Δ*hcp* mutant was generated as previously described^33^. Complementation was achieved by chromosomal integration of *hcp* together with its native regulatory region using a modified pNBU2 vector^34^. Correct integration was verified by PCR and genome sequencing. Detailed strain construction procedures are described in Supplementary Methods.

### Growth assays

Wild-type (WT), Δ*hcp*, and complemented (Δ*hcp* complemented) strains were grown in the presence or absence of nitrate or nitrite, and growth was monitored spectrophotometrically (OD600) over 24 hours.

### Human plaque collection and *ex vivo* biofilms

Supragingival plaque samples were collected from ten healthy pediatric subjects under an approved VCU IRB protocol (HM20011845). Samples were pooled and used to establish *ex vivo* oral biofilms in saliva-coated microplates with or without nitrate supplementation. Biofilms were grown under microaerophilic conditions and harvested for downstream molecular analyses.

### Long-read 16S sequencing

DNA was isolated from *ex vivo* biofilms and subjected to full-length 16S rRNA gene sequencing using Oxford Nanopore technology. Taxonomic assignment was performed against the NCBI Targeted Loci database. Detailed bioinformatic parameters are provided in Supplementary Methods.

### Nine-species oral biofilm model

A defined nine-species oral biofilm model consisting of early, intermediate, and late colonizers was established on saliva-coated hydroxyapatite surfaces. Biofilms were grown with or without nitrate supplementation and harvested for transcriptomic and quantitative analyses after seven days.

### RNA isolation and metatranscriptomics

RNA was isolated from biofilms, ribosomal RNA-depleted libraries were prepared, and samples were sequenced on an Illumina NextSeq 2000 platform. Reads were competitively mapped to reference genomes of all nine species. Reads were mapped using Bowtie2^35^. Read summarization was performed using FeatureCounts^36^. Differential expression analyses were performed using DESeq2^37^. Functional enrichment analyses were conducted using ClusterProfiler^38^.

### Quantitative PCR and nitrite measurements

Species abundance in biofilms was determined by qRT-PCR using species-specific 16S rRNA gene primers. Nitrite concentrations were quantified using the Griess assay according to the manufacturer’s instructions. Primer sequences are provided in Supplementary Table S1.

### P. gingivalis and V. atypica Co-cultures

Co-cultures of *P. gingivalis* and *V. atypica* were established in artificial saliva medium supplemented with nitrate or lactate. Population dynamics were monitored over 72 h using species-specific quantitative PCR calibrated against CFU standard curves.

## Results

### Nitrite, but not nitrate, causes nitrosative stress in *P. gingivalis*

Previously we have demonstrated the importance of Hcp for growth of *P. gingivalis* in NO_2_ and NO^33^. When grown in NO_2_, a mutant form of *P. gingivalis* lacking *hcp* gene does not grow (Fig. 1). Notably, this does not extend to NO_3_, there is no difference in growth. Complementation of the *hcp* gene restores growth of *P. gingivalis* under nitrosative stress conditions.

**Figure 1.**
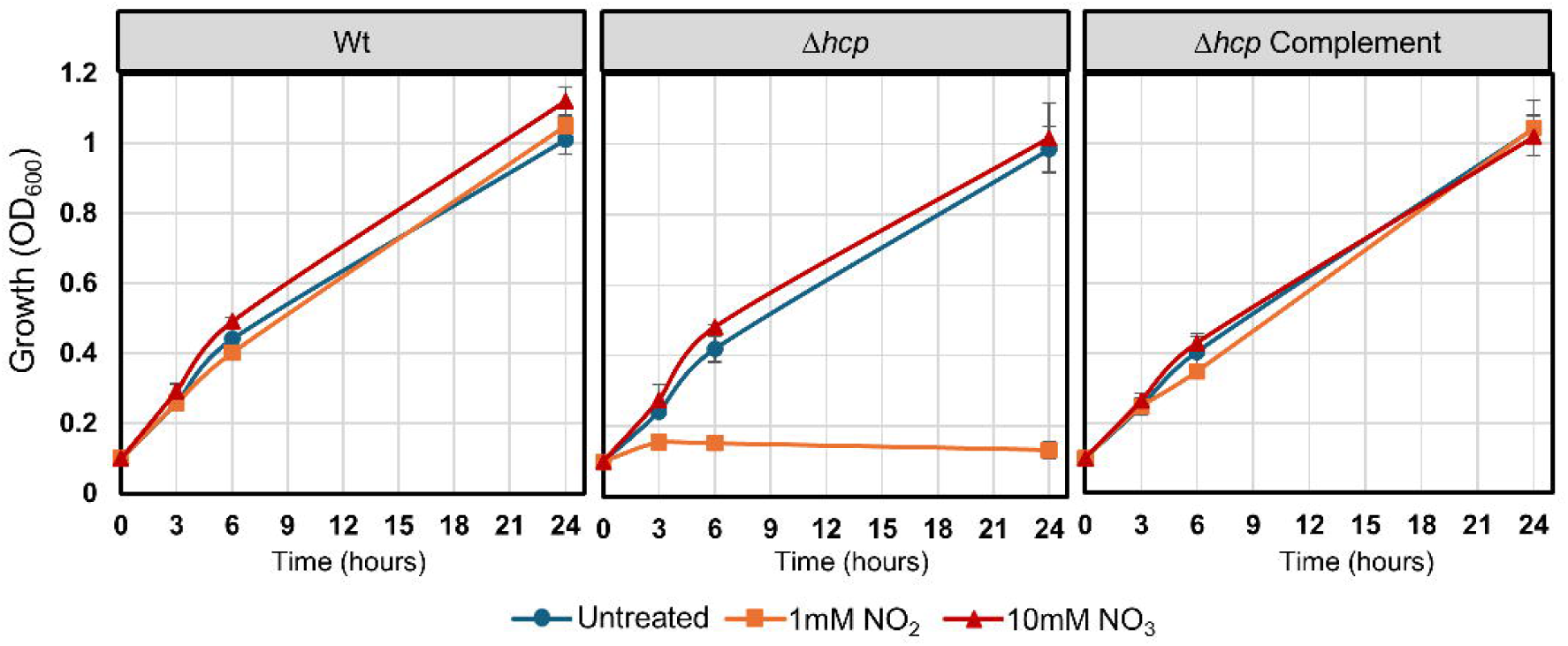
- Deletion of *hcp* sensitizes *P. gingivalis* to NO_2_ but not NO_3_. *P. gingivalis* strains were grown in mycoplasma media that was supplemented with 1mM NO_2_, 10mM NO_3_, or unsupplemented.

### Establishment of *ex vivo* plaque biofilms

To determine if NO_2_ derived from NO_3_ reducing biofilms was capable of inhibiting *P. gingivalis* growth and to determine if the Hcp reductase was important for survival in a multispecies biofilm setting, an *ex vivo* plaque biofilm model was developed. Supra-gingival plaque was taken from healthy patients (N=10) and pooled. Biofilms were initiated and spiked with or without wild type and Δ*hcp P. gingivalis* and grown with 0mM, 5mM, or 25mM NO_3_ for 24 hours in an artificial saliva media. Each biofilm growth condition was completed 3 times. The composition of the *ex vivo* plaque biofilms was determined by 16s rRNA long-read sequencing analysis.

Nitrate was supplemented in two concentrations, 5mM and 25 mM. The concentration of 5mM can be obtained in the oral cavity post nitrate rich vegetable consumption, a safe and recommendable strategy to increase nitrate levels^18,23,39,40^. The concentration of 25mM was chosen as it represents a potential topical application of probiotic nitrate.

### Community composition of biofilms is affected by nitrate

Irrespective of the presence of *P. gingivalis*, the most dominant genera at 0mM NO_3_ was *Streptococcus,* making up >50% of the total abundance. In all untreated samples the 5 most common genera were *Streptococcus, Veillonella, Gamella, Haemophilus,* and *Granulicatella* (Fig. 2A). The 16s rRNA sequencing of the untreated samples reveals that the culture conditions maintain a high degree of species diversity and are not dominated by a single species or genus and reflects their clinical origin (supragingival plaque).

**Figure 2.**
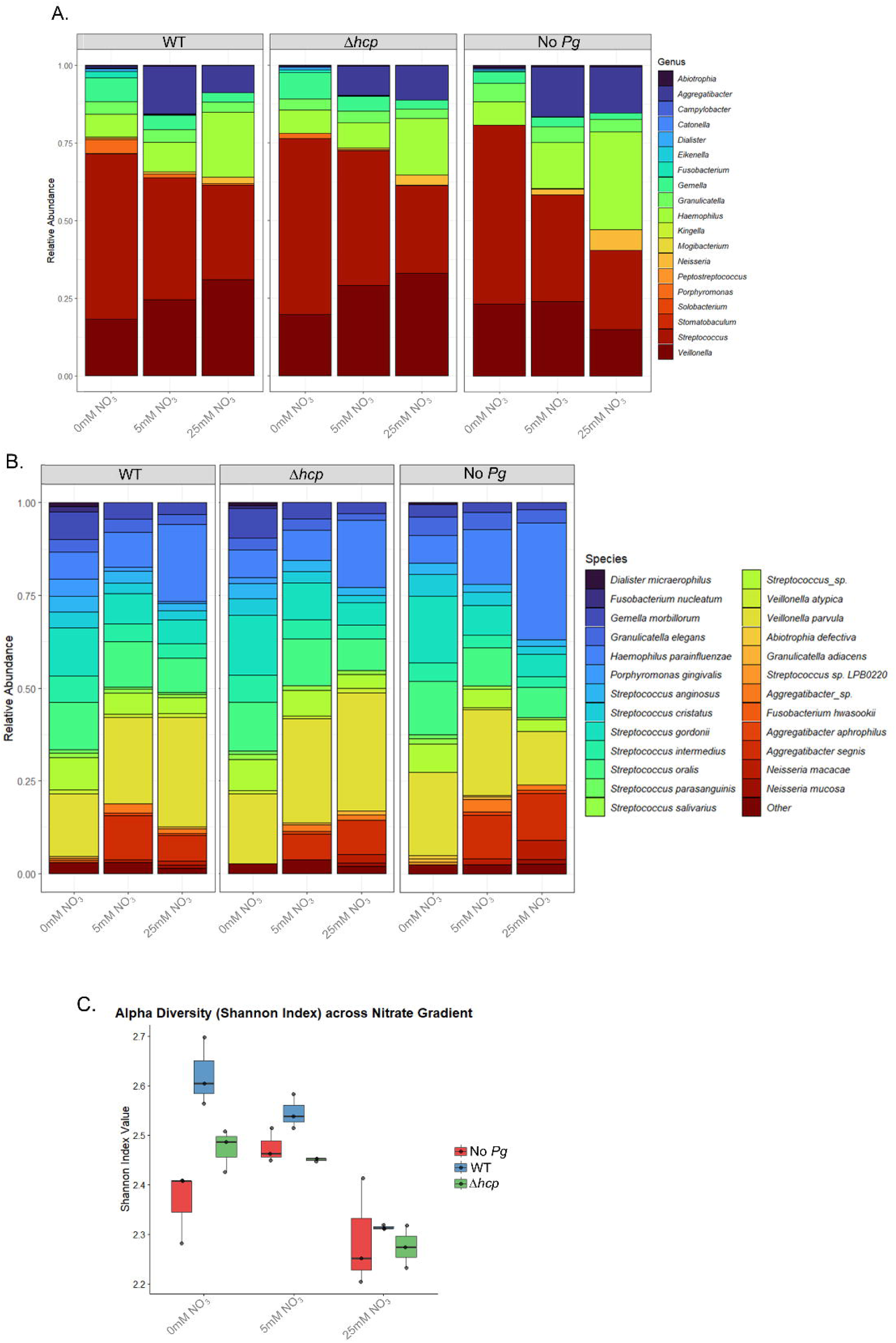
**- The effect of nitrate supplementation on *ex vivo* plaque biofilm composition.** *Ex vivo* plaque samples were grown in artificial saliva media supplemented with 5mM or 25mM NO_3_ for 24 hours. Stacked bar graphs of the relative abundance at the **A** genus level and **B** species level. Data represents averages of 3 biological replicates **C** Box and whisker plot of the alpha diversity (Shannon Index).

The relative abundance of major species in the unspiked biofilms is shown in Fig. 2B. The most dominant single species was *V. parvula*. Despite being the most dominant genera no single *Streptococcus* species dominated the biofilm. However, there was a high species diversity of *Streptococcus,* with major contributions from *S. gordonii, S. oralis, S. cristatus,* and *S. intermedius,* all health associated-*Streptococcus* species, in addition to an undefined *Streptococcus* species. Finally, a significant amount *H. parainfluenza* and *Aggregetibacter* species was present in the biofilm. This agrees with previous work, revealing a significant amount of these species in supragingival plaque from healthy sites^41^.

Nitrate treatment triggered dramatic expansions of nitrate reducing commensal species (Fig. 2B). *Haemophilus* parainfluenzae expanded from just 7.4% (0mM) to 10.8% (5mM), before becoming a dominant species at 23.4% (25mM). Well documented nitrate reducer *Veillonella parvula* also steadily increased from 19.3% (0mM) to 25.3% (25mM).

Even more pronounced was the change in *Aggregatibacter segnis* which was nearly undetectable in the absence of NO_3_ (0.07%) but increased to 10.2% (5mM) and 9.5% (25mM) Fig. 2B, Fig. 3C) Similarly, no members of the *Neisseria* genus were detected in untreated samples. However, biofilms cultivated in NO_3_ increased *Neisseria* relative abundance to 2.1 % (5mM) and 6.7% (25mM) (Fig. 2B, Fig. 3C). Using differential analysis, we found that the most significant increase was found in the species belonging to genera *Neisseria* (*N. macacae, N. mucosa*) and *Aggregatibacter (A. segnis, A. aphrophilus)* (Fig. 4A-C; Fig. S1).

**Figure 3.**
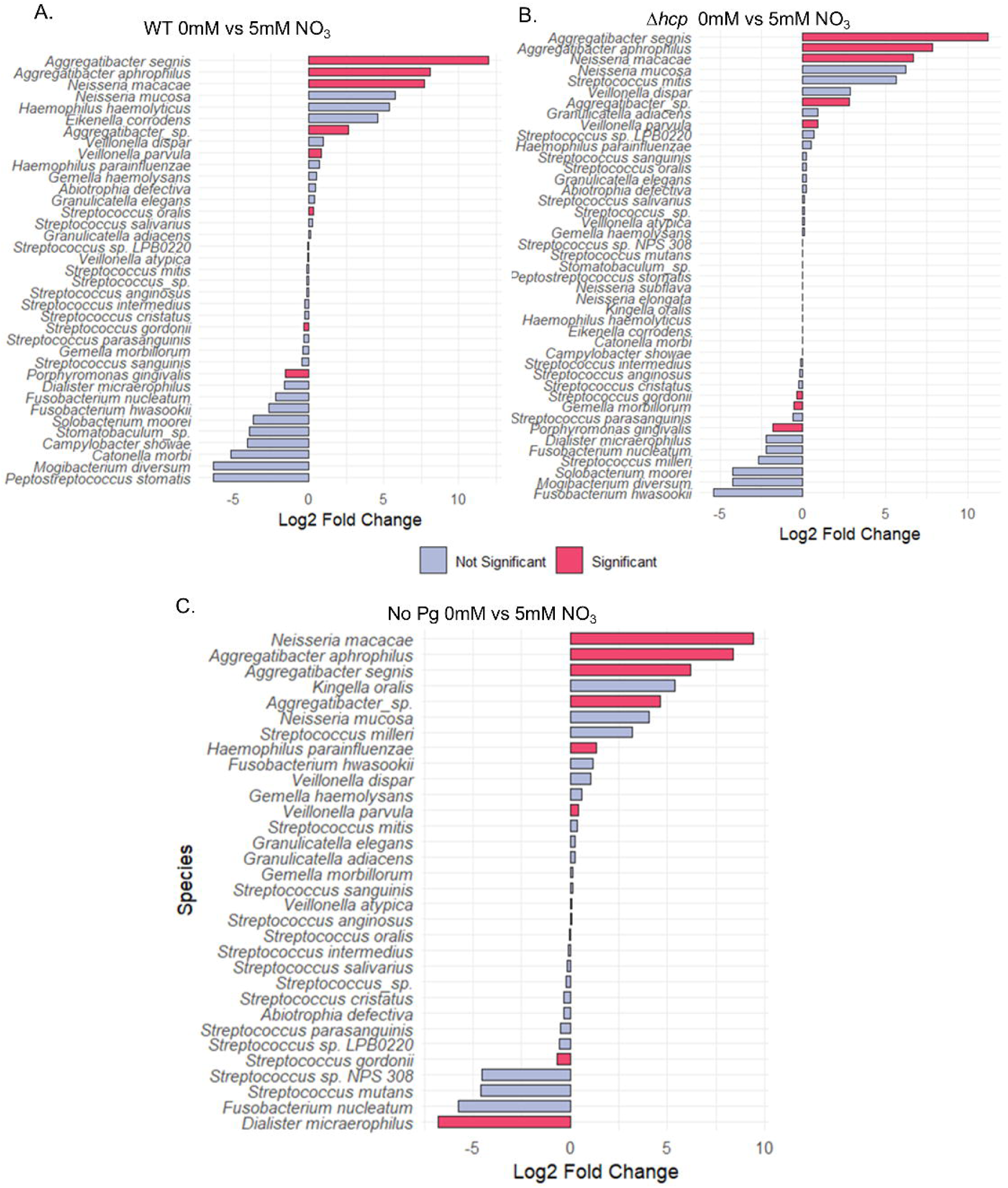
- Nitrate supplementation induces species changes in the *ex vivo* biofilms. Differential analysis of plaque biofilms grown with 0mM NO_3_ vs. 5mM NO_3_. Log2 fold change of relative abundance of all species in **A** WT *P. gingivalis* spiked biofilms, **B** Δ*hcp* spiked biofilms, and **C** unspiked biofilms. Species identified by red bars indicate significantly different changes in population (padj <0.05). Each biofilm was repeated n=3 biological replicates.

**Figure 4.**
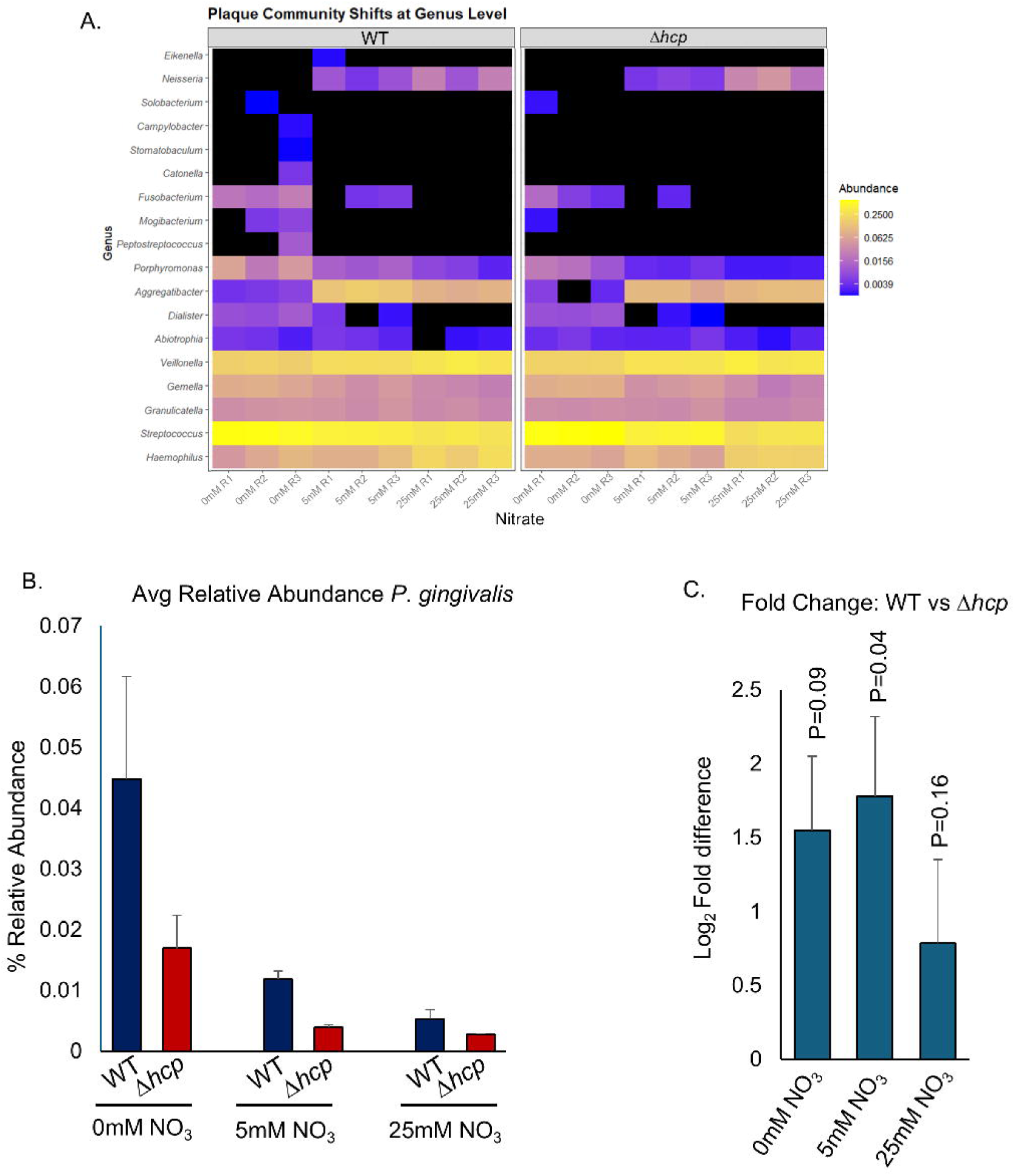
- Nitrate supplementation represses the levels of *P. gingivalis* in *ex vivo* plaque biofilms. *Ex vivo* plaque samples were grown in artificial saliva media supplemented with 5mM or 25mM NO_3_ and spiked with WT or *Δhcp P. gingivalis.* **A** Heat map of the shifts in the plaque community at the genus level. Each individual square represents a single biological replicate (n=3). **B** The average relative abundance of Wt and *Δhcp P. gingivalis* in untreated, 5mM NO_3_ and 25mM NO_3_ biofilms. **C** Log_2_ fold change in *P. gingivalis* comparing the Wt vs *Δhcp* strains at 0mM, 5mM, and 25mM NO_3_.

The expansion of nitrate reducing bacteria in the biofilm appeared to come at the expense of early colonizers. Early colonizer *Streptococcus* decreases progressively from 57.6% (0mM) to 34.3% (5mM) and 25.5% (25mM). The two most dominant species, *S. gordonii* and *S. oralis* decreased from 15.7% (0mM) to 6.16% (25mM) and 13.58% (0mM) to 8.58% (25mM) respectively. Finally, *Dialister micraerophilus* had the most significant relative decrease when grown in NO_3_ (at 5mM or 25mM). However, it was a small portion of the relative abundance in the untreated biofilm (only 0.55%) (Fig. 3C).

Crucially the pathobionts belonging to the genus *Fusobacterium* also decreased with NO_3_ supplementation. In the untreated biofilm *Fusobacterium (F. nucleatum, F. hwasookii)* had a relative abundance of 2.1%, but were eliminated in biofilms supplemented with NO_3_ (5mM <0.01%, 25mM undetected) (Fig. 2A, Fig. 3C, Fig. S1)

Comparison of the No *Pg-*spiked biofilms receiving different NO_3_ treatments indicates changes in species diversity and richness. At the genus level, increasing in NO_3_ leads to a decrease in the dominance of *Streptococcus*, and an increase in *Veillonella* and *Haemophilus* (Fig. 2A). At 5mM NO_3_ *Veillonella* is the dominant genus and at 25mM NO_3_ *Haemophilus* is the dominant genera. This corresponds with a decrease in the overall diversity of microbiomes.

### Nitrate mediated suppression of *P. gingivalis* in *ex vivo* plaque biofilms

We evaluated the abundance of *P. gingivalis* WT and Δ*hcp* spiked into plaque biofilms grown in a gradient of nitrate concentrations (0mM, 5mM, 25mM) (Fig. 4). In untreated biofilms, WT *P. gingivalis* establishes robust colonization, reaching a mean abundance of 4.5% (±1.19) (Fig. 2AB; Fig. 4AB).

The addition of nitrate resulted in a dose dependent suppression of *P. gingivalis*. The addition of 5mM NO_3_ resulted in a decrease in WT abundance to 1.18% (±0.09%), a 74% reduction from baseline. Increasing nitrate concentrations to 25mM further reduces WT abundance to 0.53% (±0.11%) (Fig. 4B).

### Hcp confers a fitness advantage in multispecies plaque biofilm environments

To determine if *hcp* protects *P. gingivalis* from RNS in *ex vivo* biofilms, we monitored the Δ*hcp* mutant strain. The Δ*hcp* strain exhibited reduced abundance compared to the WT across all culture conditions (Fig. 4AB). Even in untreated controls (0mM NO_3_) the mutant strain only achieved a relative abundance of 1.69% (±0.38%), a 69% reduction when compared to the wildtype (Fig. 4B). When treated with NO_3_, the mutant was further suppressed to 0.39% (±0.04%) at 5mM and 0.28% (0.01%) at 25mM.

Using differential analysis to compare WT and Δ*hcp*, we found that there was a statistically significant difference between the WT and Δ*hcp* at 5mM NO_3_ (adjusted P=.041) (Fig. 4C). However, at 25mM NO_3_ this trend is dampened, as the narrowing gap between WT and Δ*hcp* indicates that the RNS generated is capable of overwhelming the detoxification machinery, leading to uniform pathogen suppression.

### Addition of *P. gingivalis* changes biofilm composition

By comparing untreated biofilms spiked with WT *P. gingivalis* to unspiked controls, we identified certain interactions that *P. gingivalis* may drive. The addition of *P. gingivalis* did not dramatically affect the overall state of the biofilms, but did cause a change in the overall alpha diversity (Fig. 2C). In the absence of *P. gingivalis*, *F. nucleatum* remained constricted at low abundances (0.27%). However, spiking with *P. gingivalis* promoted a 5.4-fold increase in *F. nucleatum* (1.46%) (Fig. S2, Fig 2A). Notably, this synergistic effect was attenuated in the *Δhcp* spiked biofilms, with *F. nucleatum* only reaching 0.69%. Conversely, WT *P. gingivalis* presence actively suppressed dominant healthy oral commensals; *Veillonella parvula* dropped from 22.42% to 16.83%, and *Streptococcus gordonii* dropped from 17.94% to 13.00% upon wild-type introduction (Fig. 2A; Fig. S2).

The changes in biofilm composition can be summed up in the diversity index of the biofilms. In unspiked biofilms, the overall alpha diversity (Shannon Index) was significantly increased when supplemented with 5mM NO_3_ (Fig. 2C). However, at 25mM NO_3_ the alpha diversity was decreased as it selects for the nitrate reducing species and species that can survive with elevated nitrosative stress. Expectedly, the *P. gingivalis* spiked samples have a higher alpha diversity due to the addition of a significant amount of *P. gingivalis* to the biofilms but the increase in other pathobionts (such as *Fusobacterium*) contributes to this as well. As NO_2_ is added, the diversity of the biofilms begins to coalesce, as *P. gingivalis* and other nitrosative stress sensitive species are selected against, a uniformity in the diversity emerges.

### The effect of nitrate on a nine- species biofilm model

To verify results from our *ex vivo* plaque sequencing and further explore mechanistic insights of nitrosative stress response, a multi-species *in vitro* biofilm model containing a 9-species of oral bacteria was developed. Biofilms were grown on saliva coated hydroxyapeptite pegs and seeded in a sequential manner: (1) Early colonizers (*S. oralis, S. gordonii, C. matruchotii*) were added to establish adherence; (2) Bridging species (*F. nucleatum, V. atypica, N. perflava*) were added next to adhere to early colonizers; (3) finally, late colonizers (*P. gingivalis, P. intermedia, C. sputigina*), were added last along with with NO_3_ to determine if NO_3_ represses colonization (Fig. 5A). By forming biofilms in this way, we mimic the binding of early colonizers to the salivary pellicle followed by the binding and colonization of bridging in late colonizers in dental plaque. Biofilms were grown for 7 days under microaerophilic conditions (6% O_2_). Under these conditions, we found that the peg adhered biofilm was consistently reproducible when cultured with or without nitrate.

To ensure nitrate reduction was occurring, a sample of media was taken 3 hours post media exchange and we found NO_2_ production had spiked (Fig. 5B). We found the levels of NO_2_ to be 300-400 µM, indicating that nitrate reduction was occurring and there was metabolic activity in the biofilm.

RNA was extracted to assess levels of viable bacteria by qRT-PCR. We found the relative levels of *P. gingivalis* significantly lower when grown in 5mM NO_3_ compared to untreated controls (Fig. 5C). In addition, there were marked increases in the levels of nitrate reducers *V. atypica* and *C. sputigina*.

Next, we compared the Δ*hcp* strain to the WT. We found that there was a significant decrease in the amount of *P. gingivalis* present when grown in 5mM NO_3_ (Fig. 5D). Similarly, when using a lower amount of NO_3_ we were still able to see a decrease in both WT and mutant *P. gingivalis* strains, although this effect was expectedly blunted (Fig. 5D).

### Metatranscriptomics profiling of nine - species *in vitro* biofilm

Competitive meta-transcriptomics mapping was performed on rRNA depleted mRNA libraries generated from biofilms (Fig. 6). RNA was isolated from biofilms 3 hours post final media refresh (Day 7) to capture optimal transcription of metabolically active bacteria. Under baseline conditions, the levels of *N. perflava* and *C. matruchotii* transcripts were insufficient for adequate differential expression analysis (Fig. 6A). *P. gingivalis* persisted in the untreated biofilm at a low abundance (<0.5%) and we were capable of mapping a significant number of reads to the untreated samples.

**Figure 5.**
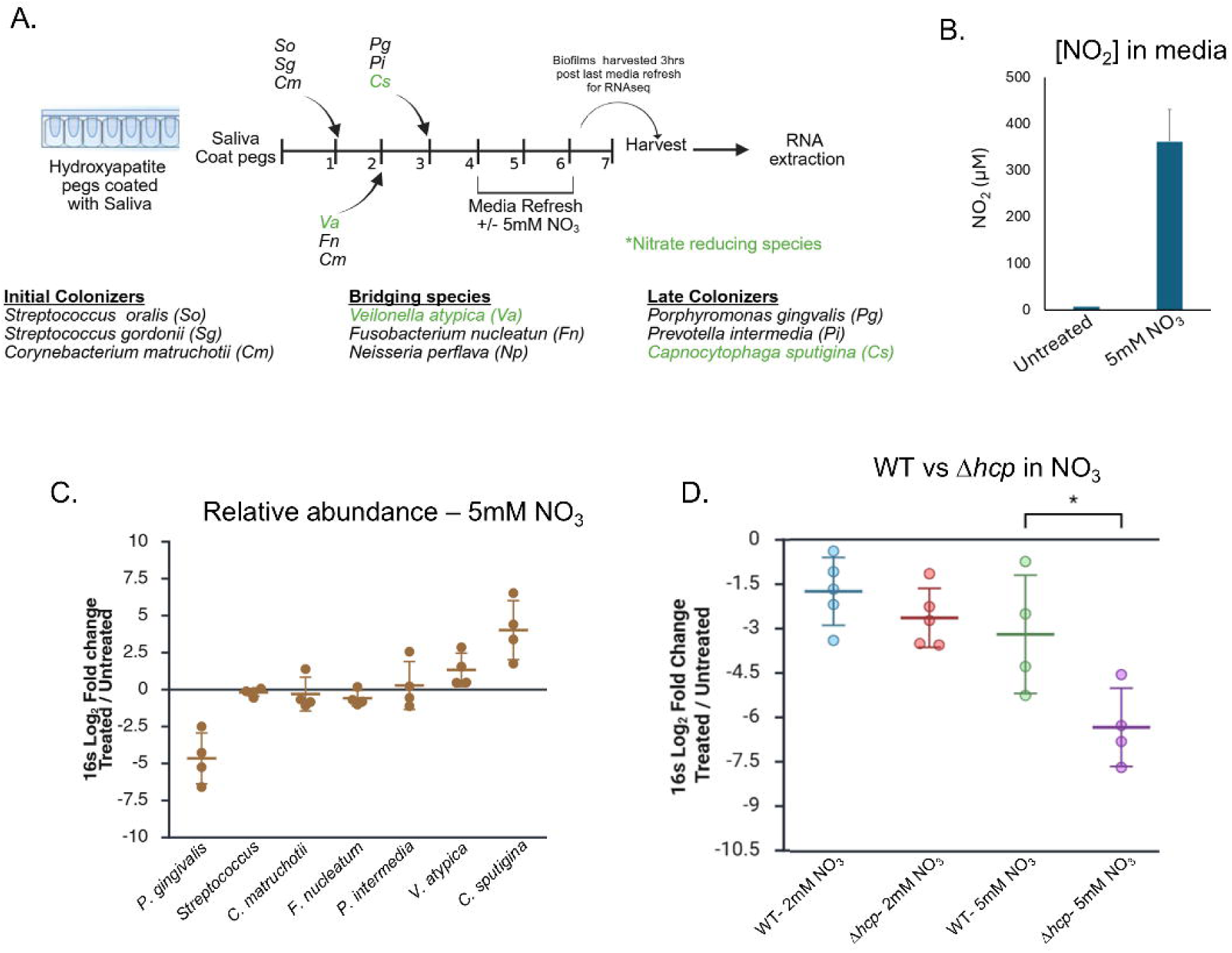
- Nitrate alters the composition of nine species *in vitro* biofilm. **A** Schematic overview of biofilm seeding and growth over 7 days. **B** Concentrations of nitrite in biofilm supernatant assessed by Griess reagent **C** qRT-PCR quantification of bacterial species comparing 5mM NO_3_ treated to untreated biofilms. Samples are representative of log_2_ fold change of each bacteria normalized to a Univ16s count. **D** The log_2_ fold change between wildtype and *Δhcp P. gingivalis* supplemented with 2mM or 5mM NO_3_.

**Figure 6.**
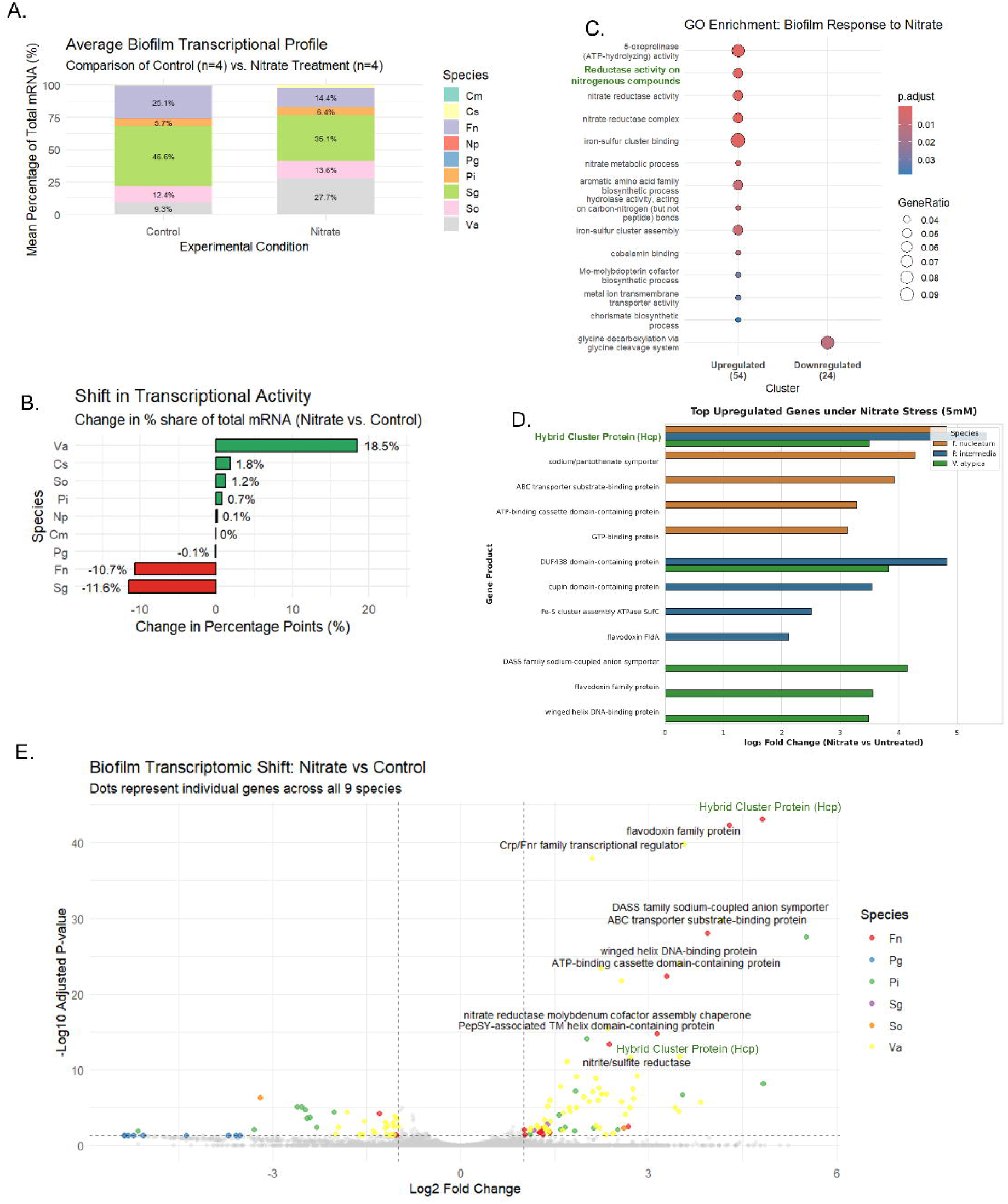
- Metatranscriptomics analysis of nitrate reducing *in vitro* biofilms. All data is representative of n=4 biological replicates. **A** Stacked bar chart depicting the mean % of total mRNA reads assigned to each species in control (0mM) and 5mM nitrate supplemented samples. **B** The change in percentage share of the total mRNA comparing cultures in control and 5mM NO_3_ biofilms. **C** Gene ontology enrichment of gene groups upregulated and downregulated in response to 5mM nitrate. **D** The top upregulated genes under nitrate supplementation in *F. nucleatum, P. intermedia,* and *V. atypica.* Bar chart expressing the log_2_ fold change of each gene or gene homologue. **E** Volcano plot of most differentially expressed genes across all 9 species in the biofilm.

In the untreated biofilm, *S. gorgonii* transcripts made up nearly half of mapped reads (46.6% of reads), followed by *F. nucleatum* (25.1% of reads) (Fig. 6A). The addition of 5mM NO_3_ to the biofilms introduced asymmetric shifts in the transcriptional abundance on the species composition, making the distribution of mapped reads more proportional (Fig. 6B). *V. atypica* emerged as the primary responder, with an +18.5 increase in total mRNA read share, reflecting its role as the primary nitrate reducer in the biofilm. Smaller transcription expansions were observed in reads of *C. sputiginia* (1.8%), *S. oralis* (+1.2%), and *P. intermedia* (+0.7%). Conversely, there were marked declines in reads of *S. gordonii* (-11.6%) and *F. nucleatum* (-10.7%) (Fig. 6B). *P. gingivalis* demonstrated a net decline of -0.1% in overall transcript share, which correlated with a near absolute population clearance, confirming the severe growth inhibition validated by qRT-PCR.

*V. atypica* showed the most robust transcriptional response to nitrate supplementation. Among the most up-regulated genes were the respiratory nitrate reductase complexes, with sub-units alpha (log_2_ FC = 2.23; padj=3.66x10^-24^), beta (log_2_ FC= 1.8; padj= 8.5x10^-10^) and gamma (log_2_ FC= 1.7; padj=7.8x10^-12^) highly expressed (Fig. 6C; Table S2). Crucially, the alpha subunit had a high baseline expression level (baseMean=21,685), signifying this pathway as a highly active metabolic influence in the biofilm.

Concurrently, *V. atypica* significantly upregulated genes involved in support of nitrate respiration involved in substrate transport and electron carriers. The DASS family sodium-coupled anion transporter (log_2_ FC=4.14) bears structural similarity to NO_3_/Na symporters^42,43^. However, the most significantly up-regulated *V. atypica* gene was a flavodoxin family protein (log_2_ FC=3.55; padj=1.35x10^-40^) (Fig. 6D-E). Flavodoxin is an iron free and oxidant resistant electron transporter, and its up-regulation is indicative of an nitrosative stress resistant change in *V. atypica* metabolism.

### Conserved nitrosative stress response across oral anaerobes

A shared, high-level upregulation of *hcp* (hydroxylamine reductase) homologs was identified among three distinct genera (Fig. 6D). In *P. intermedia, hcp* was the most highly upregulated gene in the entire metatranscriptome (log_2_ FC = 5.5; padj=2.67x10^-26^). In *F. nucleatum,* its respective *hcp* homolog was its most significantly up-regulated gene (log_2_ FC= 4.82; padj= 6.53x10^-44^). In *V. atypica,* its *hcp* homolog was similarly one of its most upregulated genes (log_2_ FC= 3.49; padj=2.14x10^-12^). This is reflected in the gene ontology (GO) analysis of the overall biofilm, with *hcp* ranking the second highest enriched group (Fig. 6C). This uniform upregulation of *hcp* serves as evidence of the community wide stress response, as these organisms actively detoxify NO and other reactive nitrogen species intermediates in the biofilm environment.

Beyond detoxification by *hcp, P. intermedia* successfully stabilizes its niche in the biofilm (+0.7% read share) by activating targeted protein repair mechanisms, specifically the entire SUF (sulfur assimilation) machinery responsible for iron sulfur cluster assembly and repair: SusC (log_2_ FC= 2.5), SusD (log_2_ FC= 1.83), and SusB (log_2_ FC= 1.56) (Fig. 6D; Table S3). In addition, *P. intermedia* also up-regulated its own flavodoxin protein (log_2_ FC= 2.11) as an iron free electron transporter.

While *F. nucleatum* robustly up-regulates *hcp,* it suffered a severe (-10.7%) loss in overall transcript share, indicating that the nitrosative stress partially outpaced its cellular defense. In tandem with its nitrosative stress response, *F. nucleatum* significantly upregulated a number of ATP dependent transporters, potentially in an attempt to balance the levels of intracellular NO_2_ or reactive nitrogen species (Fig. 6D). In addition the critical virulence factor adhesion protein *fadA* was significantly upregulated(log_2_ FC= 1.02; padj=0.042) (Table S4). This suggests a compensatory mechanism, where the microorganism increases physical adherence to the biofilm matrix to prevent detachment during metabolic stress.

The primary commensal *S. gordonii* experienced a sharp -11.6% drop in transcript share, a result that agrees with the loss in abundance seen in plaque biofilm supplemented with NO_3_. Its focused transcriptomic adaptation involved up-regulation of a formate/nitrite transporter family protein (log_2_ FC= 1.38; padj=0.0018), highlighting active attempts to pump out or regulate accumulating intracellular NO_2_ (Table S5). Similarly, *S. oralis* up-regulated an ABC transporter permease (log_2_ FC= 2.59; padj=0.0049) to potentially actively pump out excess intracellular NO_2_ (Table S6).

In our biofilm, *C. sputigina* transcript read increased (+1.8%), however, there were 0 differential regulated genes detected in NO_3_ supplemented biofilms (Table S6). The species appeared unbothered by the elevated nitrosative stress and appears to be transcriptionally unaffected by nitrate supplementation.

In contrast, *P. gingivalis* exhibited a state of complete transcriptomic collapse (Table S7). Transcripts were suppressed to very low baseline levels (baseMean <25). Even core metabolic and structural pathways lacked sufficient reads in nitrate treated samples, including glucosamine-6-phosphate deaminase and multiple proteases crucial for *P. gingivalis’* asaccharolytic metabolism.

### Nitrate disrupts Veillonella spp. - P. gingivalis synergy

We attempted to remove nitrate reducing bacteria from our nine species biofilm to determine if nitrate mediated repression of *P. gingivalis* could be reversed. However, when removing *V. atypica* from the biofilm we found that we could not effectively incorporate *P. gingivalis* in the biofilm (Fig. S3). Thus it is likely that the relationship between *P. gingivalis* and *Veillonella* helps *P. gingivalis* persist in our biofilm model^44^. Removal of *V. atypica* from the biofilm appears to lead to the loss of *P. gingivalis* (despite the presence of other species such as *F. nucleatum* and *S. gordonii,* both shown to adhere to and colonize with *P. gingivalis*).

We leveraged this *Veillonella*-*P. gingivalis* relationship by co-culturing both bacteria at very low seeding densities and observing the effects nitrate has on the growth of *P. gingivalis*. While mostly defined by its role in consumption of lactate, *Veillonella* species are capable nitrate reducers, and can utilize either lactate or nitrate respiration^13,45^. In our media conditions, either lactate or nitrate was required for *V. atypica* growth (Fig. S4A). Thus we compared co-cultures supplemented with either 10mM lactate or 10mM NO_3_ to observe the effects NO_3_ has on *P. gingivalis* growth with *V. atypica*.

We found that *P. gingivalis* does not grow effectively by itself when seeded at low cell densities, but the presence of *V. atypica* helps to overcome this limitation (Fig. 7A). In lactate co-cultures, *P. gingivalis* robustly grows from low seeding densities (∼100 CFU) over the course of 3 days (Fig. 7A). Conversely, in nitrate co-cultures, despite an initial growth over the first 24 hours, *P. gingivalis* does not surpass the growth of *P. gingivalis* mono-cultures after 72 hours. This coincides with a dramatic increase in NO_2_ concentrations, especially over 24 to 48 hours, indicating that the RNS generated by *V. atypica* arrests growth of *P. gingivalis* (Fig. 7B).

**Figure 7.**
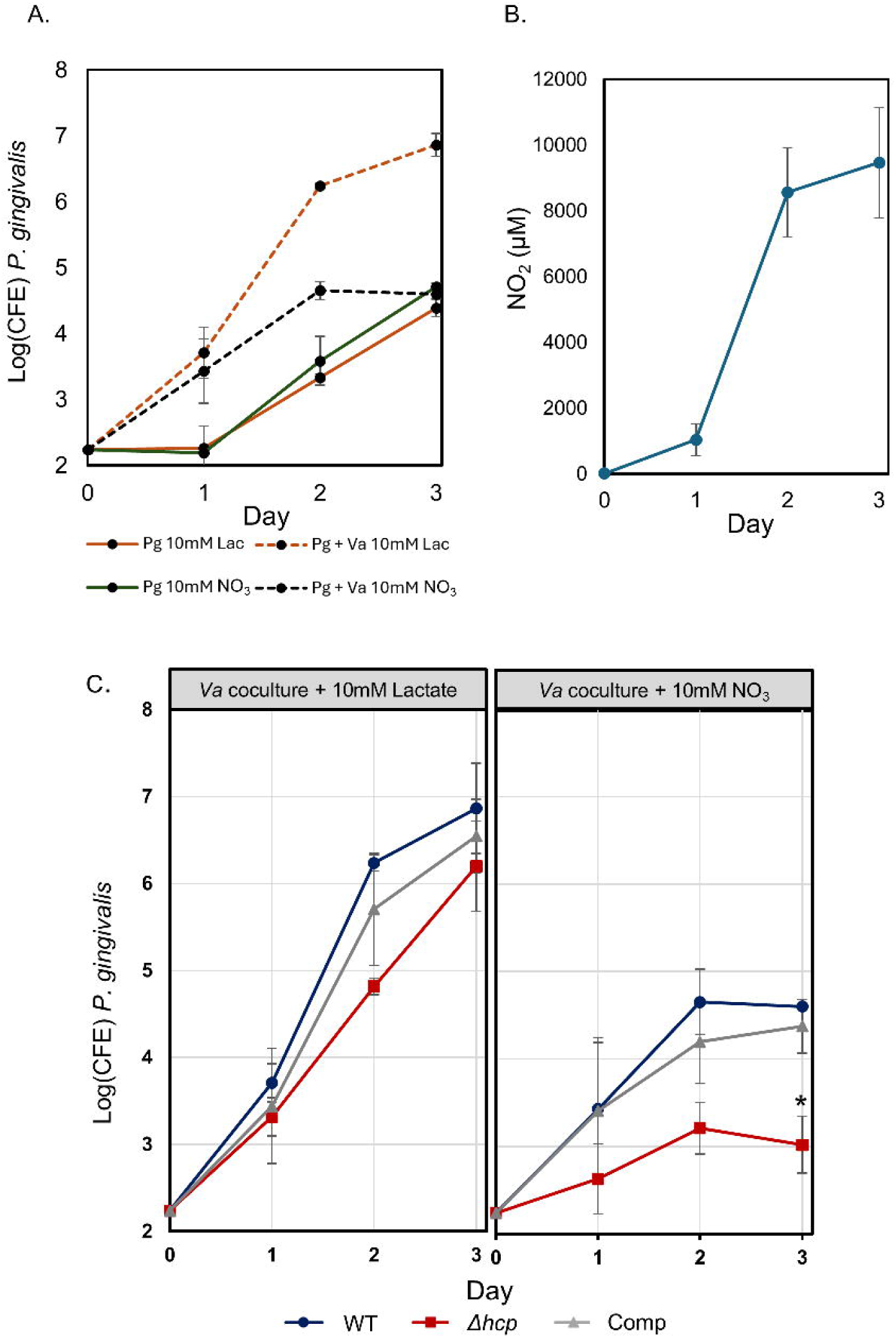
- Co-cultures of *P. gingivalis* and *V. atypica* supplemented with lactate or NO_3_. **A** Growth curve of wildtype *P. gingivalis* grown alone in artificial saliva media supplemented with 10mM lactate or 10mM NO_3_ or co-cultured with *V. atypica* with 10mM lactate or 10mM NO_3._ CFE = colony forming estimates estimated from qPCR of the 16s gene. Data is representative of the mean of 3 biological replicates. **B** Nitrite concentrations in NO_3_ supplemented *P. gingivalis-V. atypica* co-cultures over 3 days. **C** WT, *Δhcp,* or complement *P. gingivalis* co-cultured with *V. atypica* in 10mM lactate or 10mM NO_3_. Data is representative of the mean of 3 biological replicates.

In co-cultures grown with the Δ*hcp P. gingivalis* revealed near complete growth repression when grown with 10mM NO_3_ (Fig. 7C). There was no significant difference in growth after 72 hours between wildtype and Δ*hcp* strains when grown in lactate supplemented cultures. Complementing the Δ*hcp* mutant strain restored the growth to wildtype levels (Fig. 7C). In the absence of *V. atypica* there was no effective difference in growth of the Wt and Δ*hcp* strains in NO_3_ media (Fig. S4B). These results highlight the capability of nitrate supplementation to disrupt the *P. gingivalis-Veillonella* synergy and the importance of *hcp* for survival and persistence in nitrate reducing environments.

## Discussion

Over the past decade, dietary nitrate (NO₃⁻) supplementation has emerged as a promising strategy for promoting oral health and host-microbiome homeostasis. Here, we demonstrate that nitrate-reducing commensals generate sufficient nitrosative stress to suppress susceptible pathogens within complex oral biofilms. Using human *ex vivo* plaque communities, we show that microbial nitrate reduction limits colonization by *P. gingivalis*. To define the underlying mechanisms, we developed a sequentially assembled nine-species biofilm model that recapitulates oral plaque development and enables controlled investigation of nitrate-dependent microbial interactions. Collectively, our findings identify nitrosative stress as a major ecological force within nitrate-reducing biofilms and establish Hcp as a critical determinant of pathogen fitness in this environment.

A major strength of this study is the development of a defined nitrate-reducing oral biofilm model. Numerous multispecies oral biofilm systems have been described, ranging from simple co-cultures to highly complex consortia designed to mimic dental plaque^46–48^. However, few models incorporate the metabolic capacity for community-wide nitrate reduction, despite increasing recognition of the importance of the nitrate-nitrite-NO pathway in oral health. By combining early, bridging, and late colonizers with nitrate-reducing commensals, our model captures key ecological features of developing plaque while providing a tractable platform for mechanistic studies of nitrate metabolism and nitrosative stress. Importantly, nitrate supplementation resulted in measurable nitrite accumulation and selective suppression of *P. gingivalis*, demonstrating that commensal nitrate reduction can generate biologically significant nitrosative stress within polymicrobial communities.

Despite growing interest in nitrate-based therapies, the molecular mechanisms by which nitrate reduction suppresses oral pathobionts remain poorly understood. Our findings identify Hcp as a central component of this process. Hcp has been characterized as a high-affinity nitric oxide reductase in several anaerobic bacteria and is increasingly recognized as a major defense against nitrosative stress^31–33,49,50^. In the present study, deletion of *hcp* dramatically impaired the ability of *P. gingivalis* to persist within nitrate-reducing biofilms. Even in the absence of exogenous nitrate, the mutant exhibited a substantial fitness defect, and nitrate supplementation reduced survival to near-undetectable levels. These observations are consistent with our previous work demonstrating the inability of the Δ*hcp* mutant to grow at nitrite concentrations exceeding 0.2 mM^33^. Together, these studies establish Hcp as an essential adaptation that enables *P. gingivalis* to withstand physiologically relevant nitrosative stress.

Metatranscriptomic analyses further revealed that nitrate reduction is primarily driven by *V. atypica*, which expanded its metabolic activity and strongly induced expression of its respiratory nitrate reductase machinery. Increased nitrate respiration elevated extracellular nitrite concentrations and fundamentally altered the chemical environment experienced by neighboring organisms. Notably, *hcp* homologs were among the most highly induced genes in *P. intermedia*, *F. nucleatum*, and *V. atypica*, while Hcp-associated oxidoreductase activity emerged as one of the most enriched functions across the community. These findings indicate that Hcp-mediated defense is not unique to *P. gingivalis*, but rather represents a conserved adaptive response employed by phylogenetically diverse oral anaerobes exposed to nitrate-derived nitrosative stress.

Although multiple species activated the Hcp response, their outcomes differed markedly. *P. intermedia* maintained community abundance while simultaneously inducing *hcp* and genes involved in Fe-S cluster biogenesis and repair, suggesting effective adaptation to nitrosative stress. Because nitrite and NO damage iron-sulfur proteins through nitrosylation and cluster disruption, coordinated activation of detoxification and repair pathways likely contributes to resilience in this species^51^. In contrast, *F. nucleatum* exhibited strong *hcp* induction yet declined in abundance, indicating that nitrosative stress partially exceeded its protective capacity. The simultaneous upregulation of adhesins and ATP-dependent transport systems suggests an alternative survival strategy centered on maintaining biofilm attachment during physiological stress.

Nitrate supplementation caused a profound collapse of the *P. gingivalis* transcriptome. This change indicates that *P. gingivalis* is particularly vulnerable to nitrate-derived nitrosative stress generated by *Veillonella* and is unable to sustain the energetic demands of both detoxification and competition within a nitrate-reducing community. The severe phenotype of the Δ*hcp* mutant further underscores the dependence of *P. gingivalis* on this pathway for survival.

A particularly notable finding was the effect of nitrate on interactions between *Veillonella spp.* and *P. gingivalis*. Previous studies have established that *Veillonella spp.* promotes the growth and persistence of *P. gingivalis* within oral biofilms^44,52^. Consistent with these observations, removal of *V. atypica* substantially reduced *P. gingivalis* abundance in our model. However, nitrate supplementation fundamentally altered this relationship. Rather than supporting pathogen growth, nitrate-respiring *V. atypica* generated sufficient nitrite levels to suppress *P. gingivalis*, effectively transforming a mutualistic interaction into an antagonistic one. This shift was even more pronounced in the Δ*hcp* mutant, which was unable to coexist with nitrate-reducing *V. atypica*. These results illustrate how changes in microbial metabolism can reconfigure ecological interactions and reshape community structure.

In summary, our findings demonstrate that dietary nitrate functions as a microbiome-modulating prebiotic that leverages commensal metabolism to generate localized nitrosative stress. This process selectively suppresses susceptible periodontal pathogens while favoring organisms capable of withstanding nitrate-derived stress. More broadly, our data identify Hcp-mediated nitrosative stress resistance as a major determinant of bacterial fitness within nitrate-reducing oral communities and provide a mechanistic framework explaining how dietary nitrate promotes the transition from dysbiosis toward a health-associated oral microbiome.

## Acknowledgements

Sequencing data included in this study was generated at the Genomics Core facility at Virginia Commonwealth University. We would like to thank Drs. Reham Alnajjar and William Dahlke for recruiting human subjects and acquiring oral plaque samples.

## Funding

This work was supported by NIH/NIDCR grant RO1DE023304 (JPL).

## Data Availability

Raw sequencing data generated in the study (16s long read sequences and metatranscriptomic reads) have been deposited in the sequencing read archive (SRA) under Bioproject accession number: PRJNA1484659.

## Competing Interests

The authors declare no competing interests

## Supplementary Figures and Tables

Table S1 - Primers used in this study

Table S2 - List of differentially expressed genes from *V. atypica.* Data are from four biological replicates (n = 4).

Table S3 - List of differentially expressed genes from *P. intermedia.* Data are from four biological replicates (n = 4).

Table S4 - List of differentially expressed genes from *F. nucleatum.* Data are from four biological replicates (n = 4).

Table S5 - List of differentially expressed genes from *S. gordonii.* Data are from four biological replicates (n = 4).

Table S6 - List of differentially expressed genes from *S. oralis.* Data are from four biological replicates (n = 4).

Table S7 -List of differentially expressed genes from *P. gingivalis.* Data are from four biological replicates (n = 4).

---

**Figure S1.**
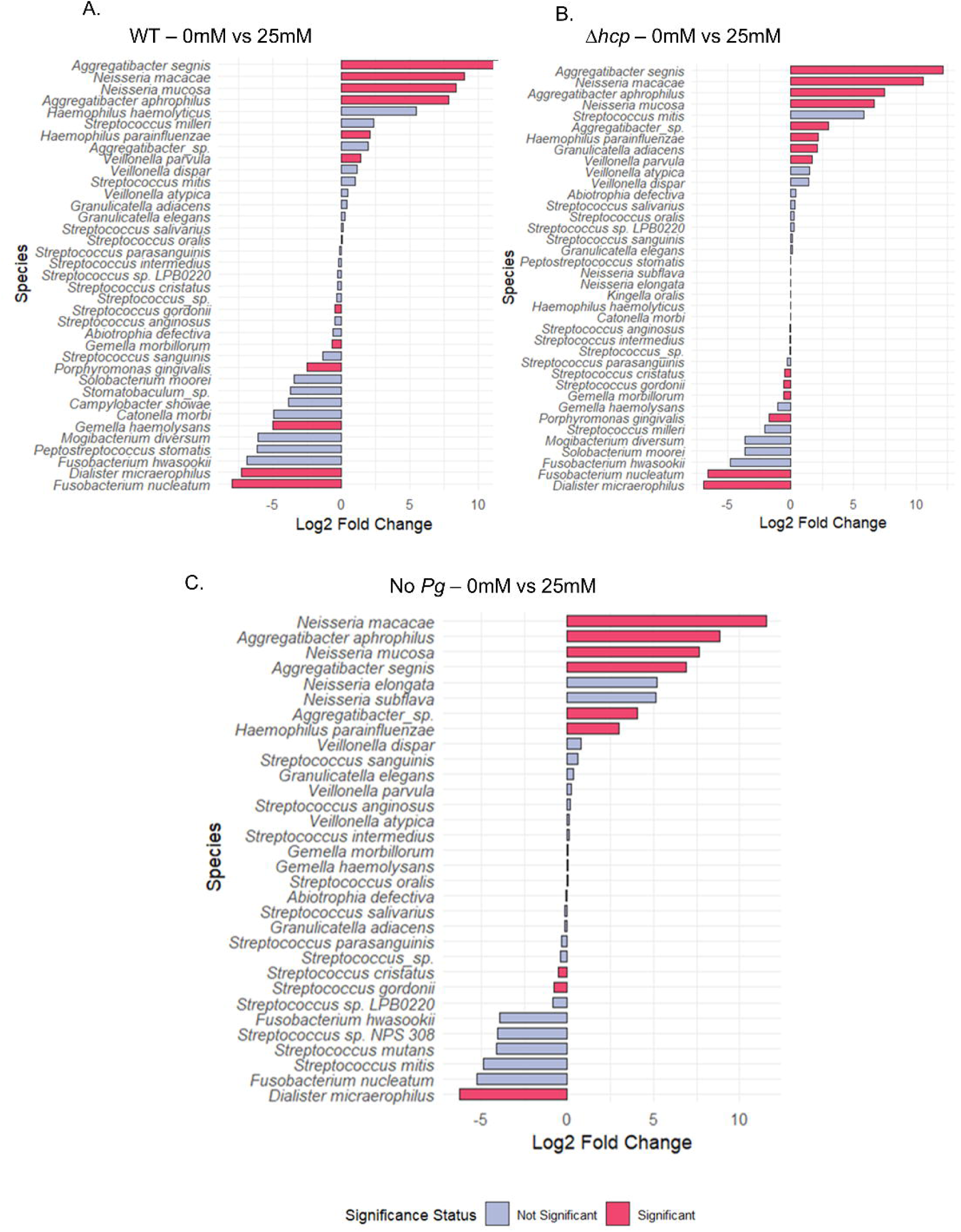
- Nitrate supplementation induces species changes in the *ex vivo* biofilms. Differential analysis of plaque biofilms grown with 0mM NO_3_ vs. 25mM NO_3_. Log2 fold change of relative abundance of all species in **A** WT *P. gingivalis* spiked biofilms, **B** Δ*hcp* spiked biofilms, and **C** unspiked biofilms. Species identified by red bars indicate significantly different changes in population (padj <0.05). Each biofilm was repeated n=3 biological replicates.

**Figure S2.**
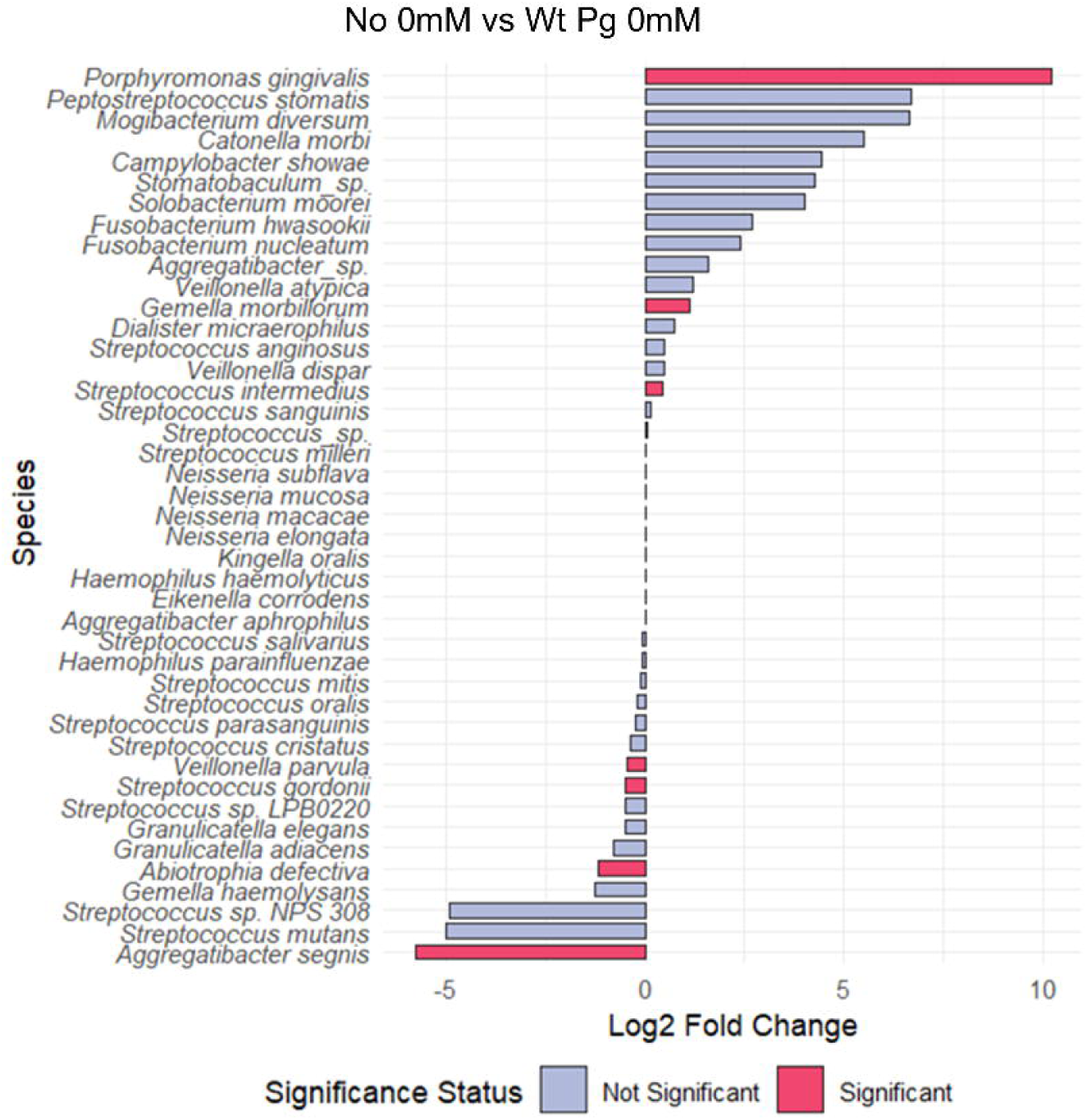
– Spiking of *P. gingivalis* WT into *ex vivo* biofilms induces species change. Differential analysis of plaque biofilms grown spiked with or without Wt *P. gingivalis* (No nitrate supplementation). Log2 fold change of relative abundance of all species. Species identified by red bars indicate significantly different changes in population (padj <0.05). Each biofilm was repeated n=3 biological replicates.

**Figure S3.**
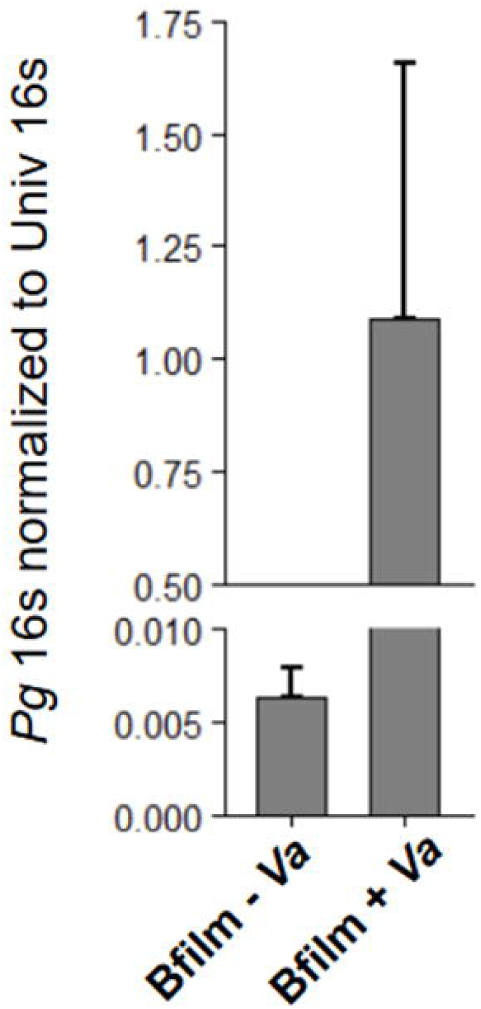
– Removal of *V. atypica* from *in vitro* biofilms inhibits *P. gingivalis* colonization in biofilms. The log_2_ fold change between wildtype and Δ*hcp* P. gingivalis supplemented with 2mM or 5mM NO_3_.

**Figure S4.**
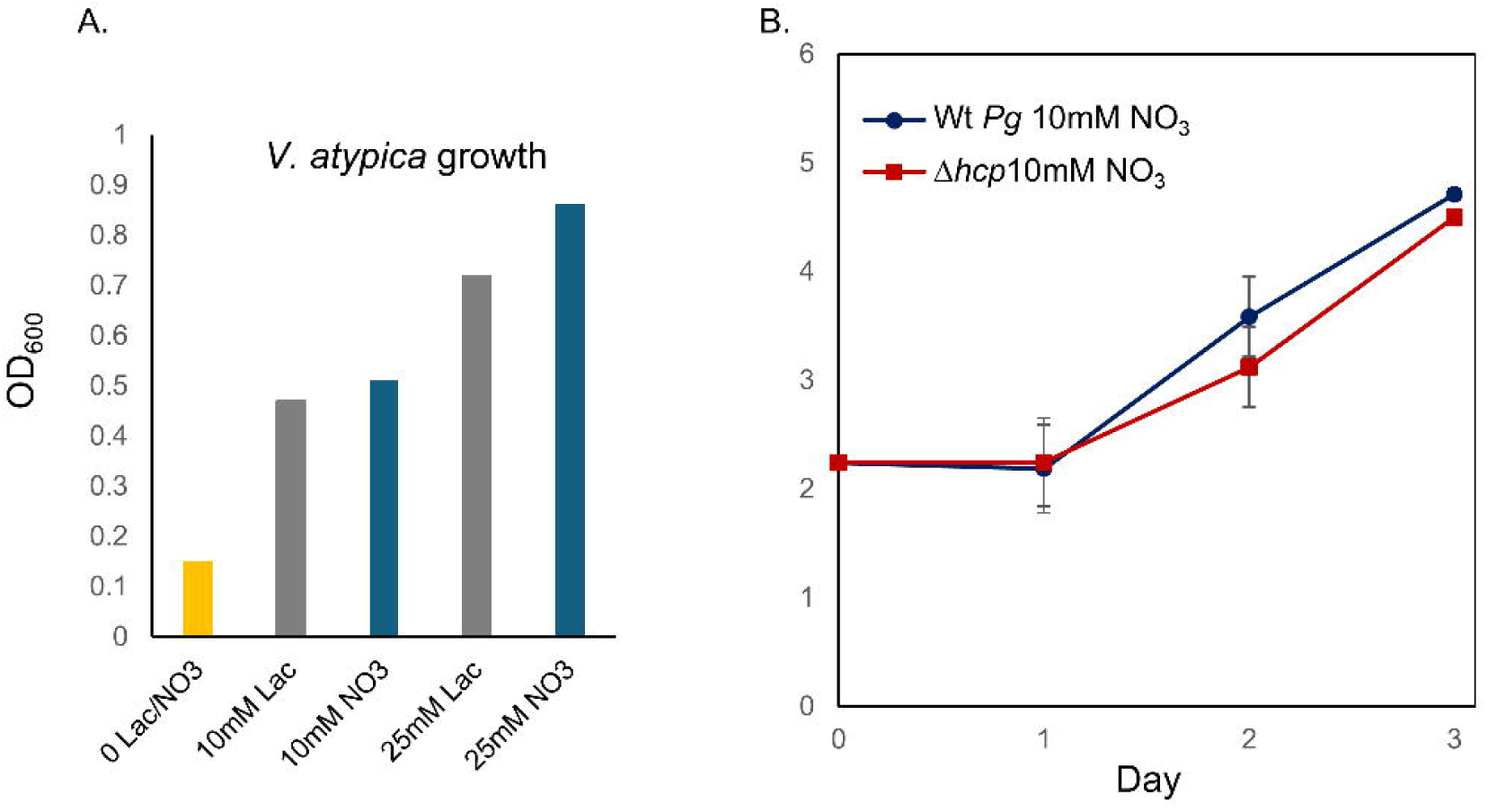
– *V. atypica* grows in lactate or NO_3_ – WT *P. gingivalis* and *Δhcp* grow similarly in NO_3_. **A** Growth of *V. atypica* in Peptone-Yeast extract media supplemented with lactate or nitrate. **B** Growth of WT and Δ*hcp P. gingivalis* in artificial saliva media supplemented with 10mM nitrate.

## Notes

### Competing Interest Statement

The authors have declared no competing interest.

