## Supplemental Materials and Methods for "Nitrate-Reducing Commensals Reshape Oral Biofilm Ecology and Reveal Hcp as a Critical Determinant of *Porphyromonas gingivalis* Persistence"

***Bacterial Strains and Growth Conditions***

Anaerobic bacteria were grown and maintained in an anaerobic environment (10% CO_2_, 10% H_2_, 80% N_2_) using a Coy Laboratories Anaerobic chamber. *Porphyromonas gingivalis* (W83)*, Prevotella intermedia* (OMA14)*, Capnocytophaga sputigina* (ATCC 33612)*,* and *Fusobacterium nucleatum* (ATCC 25586) were all maintained anaerobically on tryptic soy (TSA) blood agar plates and cultured in BHI media supplemented with 5g/L yeast extract, 5 mg/L heme and 0.1 mg/L vitamin K. *Veillonella atypica* (KON) was maintained anaerobically on BHI plates supplemented with 1.5% DL-lactate and cultured in BHI broth supplemented with lactate. *Streptococcus oralis* (ATCC 9811)*, Streptococcus gordonii* (ATCC 35105)*, Neisseria perflava* (ATCC 14799)*,* and *Corynebacterium matruchotii* (ATCC 33612) were maintained aerobically on blood agar plates and grown in supplemented BHI media. Biofilms and Co-cultures were grown in a protein rich artificial saliva medium (supplemented with FBS where noted) composed of 2 g/L proteose peptone, 1 g/L tryptose peptone, 1 g/L yeast extract, 2 g/L porcine mucin, 3.0 g/L NaCl, 1.4 g/L KCl, 1.6 mg/L ascorbic acid, 0.6 g/L KH_2_PO_4_, 5mM Urea, 9mM L-Arg, 0.1 mg/L Vitamin K, 0.5 mg/L Heme. Artificial saliva media was adjusted to pH of 7 and autoclaved.

***P. gingivalis mutation and complementation***

The *P. gingivalis Δhcp* knockout strain was made as previously described^24^. Briefly, a construct consisting of 300 bp upstream and 300 bp downstream of *hcp* coding sequence disrupted by an *ermF* cassette was assembled. The construct was PCR amplified and electroporated into *P. gingivalis* and grown overnight. The next day, bacteria were spread onto clindamycin (0.5 µg/mL) blood agar plates and grown for 6-7 days. Colonies were screened and genomes were sequenced to confirm *hcp* deletion. Complementation was achieved by using the pNBU2 vector backbone^29^. The *ermG* gene was replaced with a *tetQ* gene to infer tetracycline resistance of the complement. The whole *hcp* gene, including the upstream region containing the HcpR binding site and promoter, was then cloned into the vector. Genomic integration of the vector was achieved via conjugation using the *E. coli* S17-1 𝝀pir strain as a donor. *E. coli* containing assembled pNUB2-*tetq-hcp* plasmids were grown overnight in LB-Amp then sub-cultured into 15mLs fresh LB-Amp media. *P. gingivalis Δhcp* was grown overnight in sBHI broth in anaerobic conditions, then subcultured into 30mLs of fresh sBHI media. Both donor and recipient were grown to mid log phase (~4-5 hours). Cultures were centrifuged, washed and combined at a ratio of 2:1 recipient to donor based on OD_600_ values then centrifuged. Pellets were resuspended in 0.1mL of media and spot plated on pre-warmed TSA-Blood agar plates. Plates were incubated aerobically at 37ºC for 1 hour, followed by overnight incubation in the anaerobic chamber at 37ºC. Resulting growth from conjugation was spread onto TSA-BA+Gent+Tet plates containing 100ug/mL Gentamicin, 0.5 µg/mL tetracycline. Colonies appeared on plates 5-6 days later. The resulting colonies were subsequently tetracycline and clindamycin/erythromicin resistant. The genomic insertion of the *hcp* gene was confirmed via PCR.

***In vitro growth curve assays***

Overnight cultures of *P. gingivalis* strains were grown in BHI. The next day, cultures were diluted to a starting OD_600_ of 0.1 in mycoplasma media. Cultures were supplemented with 10mM NO_3_, 1mM NO_2_, or unsupplemented and grown for 24 hours. Growth was assessed by measuring OD_600_ at specified time points.

***qRT-PCR***

The cDNA for qRT-PCR was generated using the High Capacity cDNA generation kit (ThermoFisher). Real time qRT-PCR was performed using SYBR-green based detection on a Quant studio 3 real-time PCR system using PowerUP Syber green MasterMix (ThermoFisher). Primers targeting the 16s gene for specific genus/species were designed to assess the levels of bacteria in the biofilm. The amount of each was normalized to the level of total bacteria by using a Univ16s primer. Primers used for qRT-PCR analysis are listed in Supplementary Table S1.

***Ex vivo plaque biofilms DNA isolation and sequencing***

Microbial DNA from *ex vivo* biofilms was isolated for sequencing using ZymoBiomics DNA miniprep kit (Zymo Research) following manufacturer protocol. Pellets from technical replicates were combined and DNA isolated from the combined pellet. Cell lysis was achieved using kit lysis buffer in combination with a bead beat protocol using Lysing Matrix B bead tubs (MP Biomedical) in a FastPrep 24 Bead Homogenizer (MP Biomedical) following manufacturer recommendations for Gram+/multi-species samples. DNA was isolated in 50µL of water and stored at - 20°C.

Microbiome long read 16s amplicon sequencing was performed by Plasmidsaurus (Louisville KY, USA) using Oxford Nanopore Technology (ONT) sequencing. Isolated and purified DNA was used to amplify the full 16s gene and amplicons were used to create long- read sequencing libraries. Sequenced reads with Qscore <10, <400bp in length, or >3000bp in length were omitted from analysis. Each read was classified against the NCBI Targeted Loci database against bacterial targets (https://www.ncbi.nlm.nih.gov/refseq/targetedloci/) to determine taxonomic classifications. Taxonomic data is compiled from raw reads to generate metrics on relative abundance and differential analysis of bacterial species.

***RNA isolation***

To isolate RNA from biofilm pellets, the RNeasy kit (Qiagen) was used with a modified lysis step. Pellets were resuspended in lysis buffer and an equal volume of acid-phenol chloroform (1mL total) was added to the samples. Samples were placed in Lysing Matrix B tubes (MP Biomedicals) and bead beat 2x (1min, Max speed) using a FastPrep 24 Bead Homogenizer (MP Biomedicals). Homogenized samples were spun down and the aqueous phase was removed and diluted in an equal volume of 100% ethanol. RNA was then purified following manufacturers protocol. After elution of RNA, samples were additionally treated with the DNA-free DNA Removal kit (Thermo-Fisher Scientific) to remove residual genomic DNA.

***Plaque sampling and Patient conditions***

The study was approved by the Virginia Commonwealth University Institutional Review Board (IRB HM20011845). Our study sample included 10 patients who presented to the VCU Graduate Pediatric Dentistry clinic for routine dental care. Inclusion criteria was medically healthy children in the primary dentition and were caries free. Guardians of children gave verbal consent and waiver of assent for participation of their children.

Two plaque samples (one anterior, one posterior) were collected from each participant. All sites were air dried, and cotton roll isolation was used. Supra-gingival plaque was gently removed from the tooth via a sterile curette and stored in 1 mL of Brain Heart Infusion (BHI) medium + 10% filtered artificial saliva in a microcentrifuge tube on ice. The sample was immediately transported into an anaerobic chamber where the sample was split into two 250 µl aliquots and diluted with 250 µl of anaerobic Shi medium to lower the oxygen level of the sample. Samples were incubated overnight in an anaerobic atmosphere at 37 ºC using a Coy anaerobic chamber (Ann Arbor, MI). After overnight incubation the aliquots were stored at -80 ºC with 10% of glycerol. Sample aliquots from patients were pooled together and used as an inoculum for *ex vivo* biofilms studies.

***Ex vivo plaque biofilm growth***

Wells of a 96-well plate were coated with sterile filtered saliva overnight and allowed to air dry. Pooled plaque samples were diluted 1:10 in a media made up of artificial saliva + 5% FBS which was supplemented with 0mM, 5mM, or 25mM NO_3_. For samples with *P. gingivalis* strains spiked in, *P. gingivalis* was added to a final concentration of 0.075 OD_600_. Cultures were then placed in a micro-aerophilic environment (6% O_2_ , 5% CO_2_, 5% H_2_, 84% N_2_) and grown for 24 hours at 37°C. Cultures were grown in technical triplicates (grown on the same plate). The *Ex vivo* biofilm growth was repeated 3 times. After incubation, supernatant media was removed and biofilms were washed with PBS. Biofilms were resuspended in PBS, spun down, and stored at -80°C for nucleic acid isolation.

***Multispecies in vitro biofilm model***

A schematic overview of the *in vitro* biofilm protocol is available in figure 5A. *In vitro* nitrate reducing biofilms were created using 9 oral species: *Streptococcus oralis* (ATCC 9811)*, Streptococcus gordonii* (ATCC 35105)*, Corynebacterium matruchotii* (ATCC 14266)*, Veillonella atypica (strain KON), Fusobacterium nucleatum* (ATCC 25586)*, Neisseria perflava* (ATCC 14799)*, Porphyromonas gingivalis W83, Prevotella intermedia OMA14,* and *Capnocytophaga sputigina* (ATCC 33612)*.* Biofilms were initiated and grown on hydroxyapatite coated pegs using MBEC assay biofilm inoculators (Innovotech). Before inoculation, pegs were coated with sterile filtered saliva overnight. Bacteria were added sequentially to pegs in groups of 3 (early, intermediate, and late colonizers) by removing the top (with pegs attached) and placing in a new 96-well plate. Equal amounts of each strain were added to a final OD_600_ of 0.1 in media. First, early colonizers were added to pegs (*S. oralis, S. gordonii, C. matruchotii)* in artificial saliva media supplemented with 20% FBS. Plates were grown for 24hours at 37°C in 6% O_2_ atmosphere. The next day, intermediate colonizers were added to pegs (*F. nucleatum, V. atypica, N. perflava*) in fresh AS-FBS media. Plates were grown for 24 hours at 37°C in 6% O_2_ atmosphere. Finally, late colonizers (*P. gingivalis, P. intermedia, C. sputigina)* were added to pegs in AS media (no FBS) supplemented +/– nitrate and grown for 24 hours at 37°C in 6% O_2_ atmosphere. The media was refreshed every 24 hours. After 7 days since biofilm initiation, samples were harvested by removing pegs from the top using flame sterilized pliers. Samples were placed in tubes with PBS and biofilms were removed from pegs using sonication. Pegs were discarded and cultures were spun down and pellets stored at -80°C for RNA extraction.

***Biofilm Transcriptomics***

Ribosomal-depleted RNAseq library generation of biofilms was created using the Universal Prokaryotic RNA-seq kit with Prokaryotic AnyDeplete (NuGEN). Libraries were generated following the manufacturer's protocols. Libraries were sequenced by the VCU nucleic acid sequencing core using NextSeq2000 Illumina sequencer using a P1 flow cell and 150bp dual end reads. Sequenced reads were competitively mapped to all 9 species genomes using Bowtie2^33^. For each species, read summarization was performed using featureCounts and differential expression analysis was performed using DESeq2^34,35^. ClusterProfiler was used for gene ontology enrichment analysis^36^.

***Determination of nitrite concentrations***

The levels of nitrite in cell culture media were assessed using the Griess Reagent Nitrite Measurement Kit (Cell Signaling Technology). Cell culture supernates were spun down and supernatants were diluted using distilled water. Measurements were acquired by following manufacturer's protocol.

***Growth of P. gingivalis and V. atypica Co-cultures***

The OD_600_ of overnight cultures of *P. gingivalis* and *V. atypica* were measured and used to seed cultures in artificial saliva media to ~100 CFUs (LogCFU ≈ 2.0). Cultures were supplemented with either 10mM Na-Lactate or 10mM NaNO_3_. A sample was taken immediately after culture initiation as the D0 point. After 24 (Day 1), 48 (Day 2), and 72 hours (Day 3) an aliquot of the culture was taken and stored at -20°C. To estimate culture CFUs, culture aliquots were boiled for 10 minutes to lyse cells before being used as a template in qPCR reaction using targeted 16s primers. A standard curve was created by serial diluting cultures and plating to obtain CFU values and observe their corresponding Ct values. This standard curve was used to determine the estimated CFUs of co-culture lysates.
